# Introducing CELLBLOKS^®^: a novel organ-on-a-chip platform allowing a plug-and-play approach towards building organotypic models

**DOI:** 10.1101/2022.04.05.487165

**Authors:** Valon Llabjani, M.R. Siddique, Anaïs Makos, Afaf Abozoid, Valmira Hoti, Francis L Martin, Imran I. Patel, Ahtasham Raza

## Abstract

Human organs are structurally and functionally complex systems. Their function is driven by interactions between many specialised cell types, which is difficult to unravel on a standard petri-dish format. Conventional “petri-dish” approaches to culturing cells are static and self-limiting. However, current organ-on-a-chip technologies are difficult to use, have a limited throughput and lack compatibility with standard workflow conditions. We developed CELLBLOKS^®^ as a novel “plug & play” organ-on-a-chip platform that enables straightforward creation of multiple-cell type organ specific microenvironments and demonstrate its advantages by building a liver model representative of live tissue function. CELLBLOKS^®^ allows one to systematically test and identify various cell combinations that replicate optimal hepatic relevance. The combined interactions of fibroblasts, endothelial cells and hepatocytes were analysed using hepatic biochemistry (CYP3A4 and urea), cellular proliferation and transporter activities (albumin). The results demonstrate that optimal liver functional can be achieved in cross talk co-culture combinations compared to conventional mono-culture. The optimised CELLBLOKS^®^ liver model was tested to analyse drug-induced liver toxicity using tamoxifen. The data suggests that our CELLBLOKS^®^ liver model is highly sensitive to toxic insult compared to mono-culture liver model. In summary, CELLBLOKS^®^ provides a novel cell culture technology for creating human relevant organotypic models that are easy and straightforward to establish in laboratory settings.

## Introduction

*In-vitro* models aiming to simulate real organ functions for research need to consider complex tissue microenvironment that not only includes parenchymal cells but also their interactions with surrounding cells such as endothelial cells, fibroblasts and immune cells, in addition to the extracellular matrix composites (Hanahan & Weinberg, 2011; Joyce & Pollard, 2009; Koontongkaew, 2013). In drug discovery, high rates of drug attrition in clinical trials suggest there are limitations in current prediction capabilities of present preclinical models (both *in-vitro* and *in-vivo* models). Drug-induced liver injury (DILI) continues to be the leading cause of attrition during drug development in all phases of clinical trials as well as the number one cause of post-market drug withdrawal, accounting for 20-40 % of all cases (Fung *et al*., 2001; MacDonald & Robertson, 2009; Onakpoya *et al*., 2016).

There is a general prerequisite that novel technologies need to be simple, rapid and physiological relevant to human liver, so as to provide improved predictive models for drug discovery (Lin & Khetani, 2016; Zhou *et al*., 2019). Several *in-vitro* and *in-vivo* models are used to screen for DILI-related toxicity issues. However, the main limitations with animal tests include the differences in physiological parameters (*i.e*., genetics and metabolic processes) between humans and rodents (Ballet, 2015; Lemon & Dunnett, 2005; Roth & Ganey, 2011). Out of 150 studied hepatotoxins, both rodents (primary rat) and non-rodents (*e.g*., canine) models only detected 50% of human hepatoxic events associated with these agents (Olson et al., 2000). Additionally, standard *in-vitro* models lack immune system incorporation and there is failure to account for crosstalk with other cell types, which is important in driving their hepatic relevance (Petrov *et al*., 2018).

*In-vitro* approaches used to create more relevant organ-specific functions have conventionally involved either mixing different cell types randomly, in one well, or a sandwich culture method where cell types are built in layers, consecutively, one cell type at a time. Although using the sandwich approach aims to mimic tissue architecture the method is time consuming, difficult to reproduce and labour intensive (Dunn et al., 1991). Similarly, randomly arranged co-cultures are likely to result in a highly variably output with each experimental repeat leading to uncontrolled cell attachment, aggregation and migration that can change with different cell-cell ratios (Bhatia *et al*., 1998; Li *et al*., 2014). Furthermore, analysis of each cell type separately is not an option in co-culture models.

Although, several organs-on-chips (OoCs) microfluidic-based approaches have been recently introduced as alternatives to conventional grown cells on 2-D surfaces, these models are only in their infancy; they require comprehensive characterisation before their adaptation into drug discovery pipelines (Zhou *et al*., 2019). Their adaptation is limited due to their substantial differences to conventionally used industry standard multi-well plates that have shortcomings in both handling and biological characterisations; they differ in size (micro-chip to macrowell plates), are difficult to handle, contain different surface chemistry for cell growth (*e.g*., PDMS), limited endpoint measurements and low throughput (Beckwitt *et al*., 2018; Cui & Wang, 2019; Lin & Khetani, 2016).

CELLBLOKS^®^ is a patented (GB2553074B), open-top multi-chambered organs-on-a-chip device (Figure 1). The platform is designed in a standard SBS footprint consisting of four lines of three interconnected chambers, and in each chamber is inserted a cell growth block. Each cell growth block serves as an individual block in which different cells can be seeded. Three different kinds of blocks suitable for different cell types are available and include Barrier Blocks™, Circulatory Blocks™ and Blank Blocks™. Barrier blocks™ are used to emulate barrier functions such as gastrointestinal (GI) tract and blood brain barrier (BBB), whereas Circulatory blocks™ mimic tissues in systematic circulation. Blank blocks™ are used to isolate cell compartments which are often used as control. The cell culture blocks are connected through the channels, from which flows the culture medium between them. In the CELLBLOKS^®^ platform, each cell block can be examined separately, and different cell types can be added or removed to the system any time during the study. This can be done in a nondestructive manner, enabling versatility to building optimal organ specific models and monitoring model performance in real-time.

**Figure 1.**
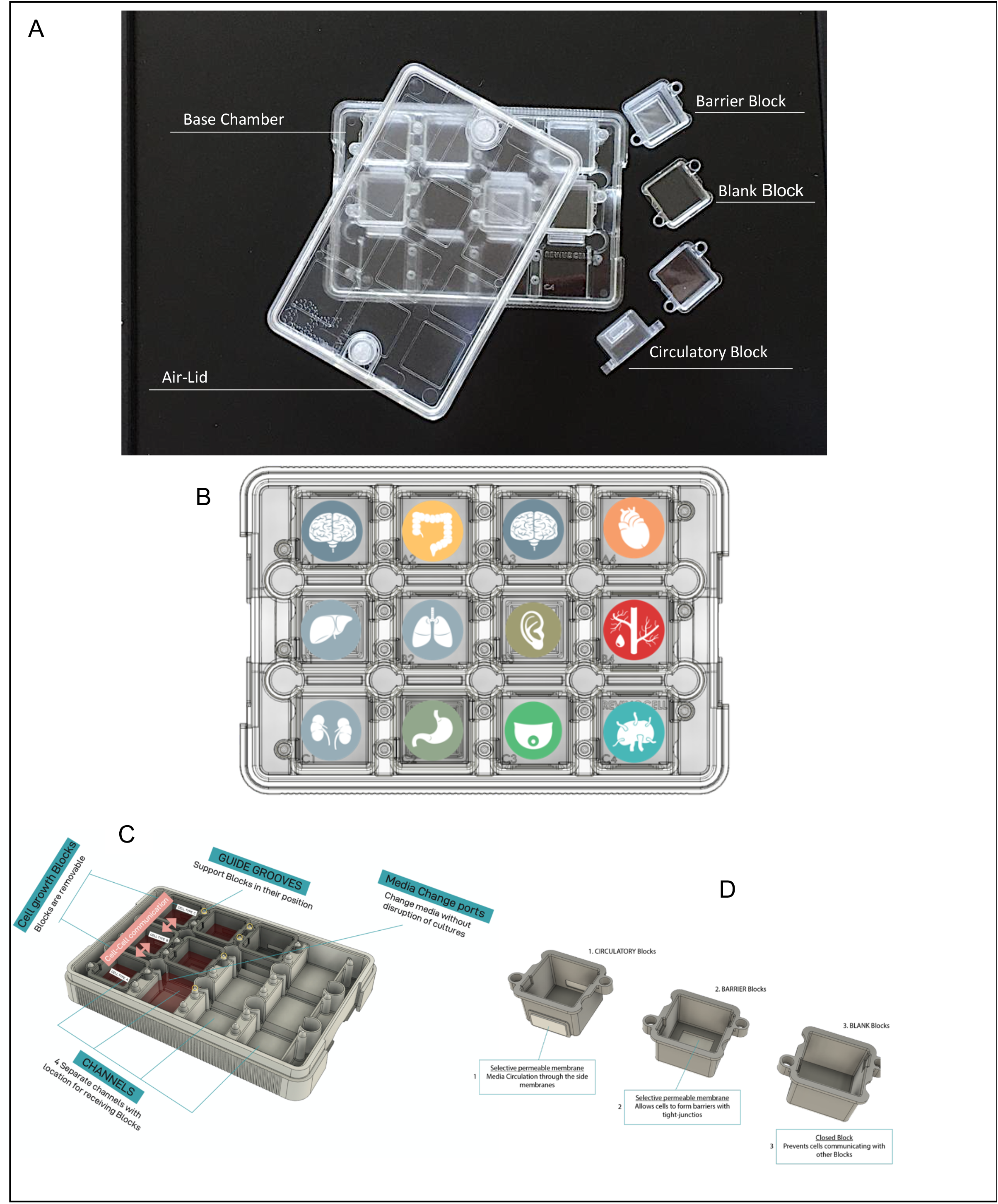
Illustration of the CELLBLOKS^®^ platform. The platform has the dimensions of a standard tissue culture well plate and is designed to allow multiple organ specific cells/tissues to grow in separate compartment blocks **(A and B)**. These are interconnected *via* cell growth blocks that maintain cells in their respective compartments but allow non-contact cell-cell communication *via* media flow channels (**C**). CELLBLOKS^®^ has four separate elongated channels with location for three separate cell growth blocks. Each channel is filled with media (3-5 ml) to allow the cell-cell communication between the blocks. Three types of blocks are used to mimic different tissue specific conditions **(D)**. ***1*. Circulatory Blocks** provide a bottom surface for cells to grow and side circulatory windows in the walls allowing selective media diffusion (both inlet and outlet, simulating organs in systematic circulation, *e.g*., liver, brain, heart, lung); ***2*. Barrier Block** that contains a selective permeable membrane on the bottom of the block, allowing cells to proliferative on a basolateral membrane (simulating epithelial cells and tissues); and, ***3*. Blank Blocks** have the same surface as the circulatory blocks for cell growth but no inlet or outlet for media diffusion. Blank blocks are used to isolate cell cultures from other compartments and are often used as controls.

Herein, we have used CELLBLOKS^®^ to seed various cell types to recapitulate liver tissue architecture in a connected interactive co-culture liver model. We hypothesise that in such settings the crosstalk between the hepatocytes, fibroblasts and endothelial cells will enhance hepato-cellular metabolic functions compared to conventionally grown hepatocyte monocultures and provide more hepatic-relevant model for drug screening. The liver model was established by using HepG2 hepatocytes, an extensively studied cell line in drug safety screening, combined with NIH/3T3 fibroblasts and human umbilical vein endothelial cells (HUVEC) that are reported to support the functions of hepatocytes in co-culture studies (Bale *et al*., 2014; Bhatia *et al*., 1998; Cho *et al*., 2010; Evenou et al., 2011; Freyer *et al*., 2017; Gebhardt *et al*., 2003; German & Madihally, 2019; Khetani *et al*., 2004; Takayama *et al*., 2013). One of the critical functions of hepatocytes in the liver is the synthesis of albumin, a protein of 585 amino acids known to play a critical role in the binding and transport of drugs, maintenance of colloid osmotic pressure, and the scavenging of free radicals (Kane et al., 2006; Ranucci et al., 2000). In this study, we used albumin production, CYP estimation and urea production to optimise CELLBLOKS^®^ liver model functions. We have also studied the toxicology profile of one of the known DILI compound, tamoxifen, to estimate the IC50 values in the CELLBLOKS^®^ liver model.

## Material and Methods

### Cell culture

Human hepatic carcinoma (HepG2) cells, human umbilical vein endothelial (HUVEC) cells and mouse fibroblast (NIH-3T3) cells were purchased from Sigma-Aldrich™. HepG2 cells and NIH-3T3 cells were cultured in T25 cell culture flasks at 37°C and 5% CO_2_ in Dulbecco’s Modified Eagle Medium (DMEM) (Gibco™) supplemented with 10% (v/v) of foetal calf serum (FCS) (Thermofisher™), 2 μM L-glutamine (Sigma-Aldrich™), 100 IU/mL penicillin (Sigma-Aldrich™) and 100 μg/mL streptomycin (Sigma-Aldrich™). HUVECs were cultured in T25 cell culture flasks at 37°C and 5% CO_2_ in an Endothelial cell Growth Media (EGM) (Cell Application™). The media was changed every 2 days and cells passaged twice weekly.

### Liver modelling customisation in CELLBLOKS^®^ platform

The human liver model is depicted using the CELLBLOKS^®^ platform (Figures 1 and 2). Hepatocytes (HepG2 cells), endothelial cells (HUVECs) and fibroblasts (NIH-3T3 cells) are seeded in tri-culture, co-culture, and mono-culture combinations to investigate various cell combinations to optimize liver function (Figure 2A). Circulatory blocks™ that allow exchange of media components between cell block compartments were selected and were inserted in chambers: [A1 - B1 - C1], [A2 - B2], [A3 + B3], [A4 + B4] to allow the testing of 2-way or 3-way co-culture set-up combinations. Blank blocks™ were inserted in chambers [C2], [C3] and [C4] to isolate cells (no media exchange) from other cell block compartments; these latter were used as mono-culture controls for each individual cell type. Cells in Circulatory blocks™ can communicate through 1μm polycarbonate selectively permeable membrane windows incorporated into each block, *via* media circulation in the interconnected chambers (Figure 2A, red arrows). In contrast, cells in Blank blocks™ are isolated from other cultures and cannot communicate with other cell types. These blocks are designed to study monoculture controls with no co-culture or flow interactions in the same experimental system.

**Figure 2.**
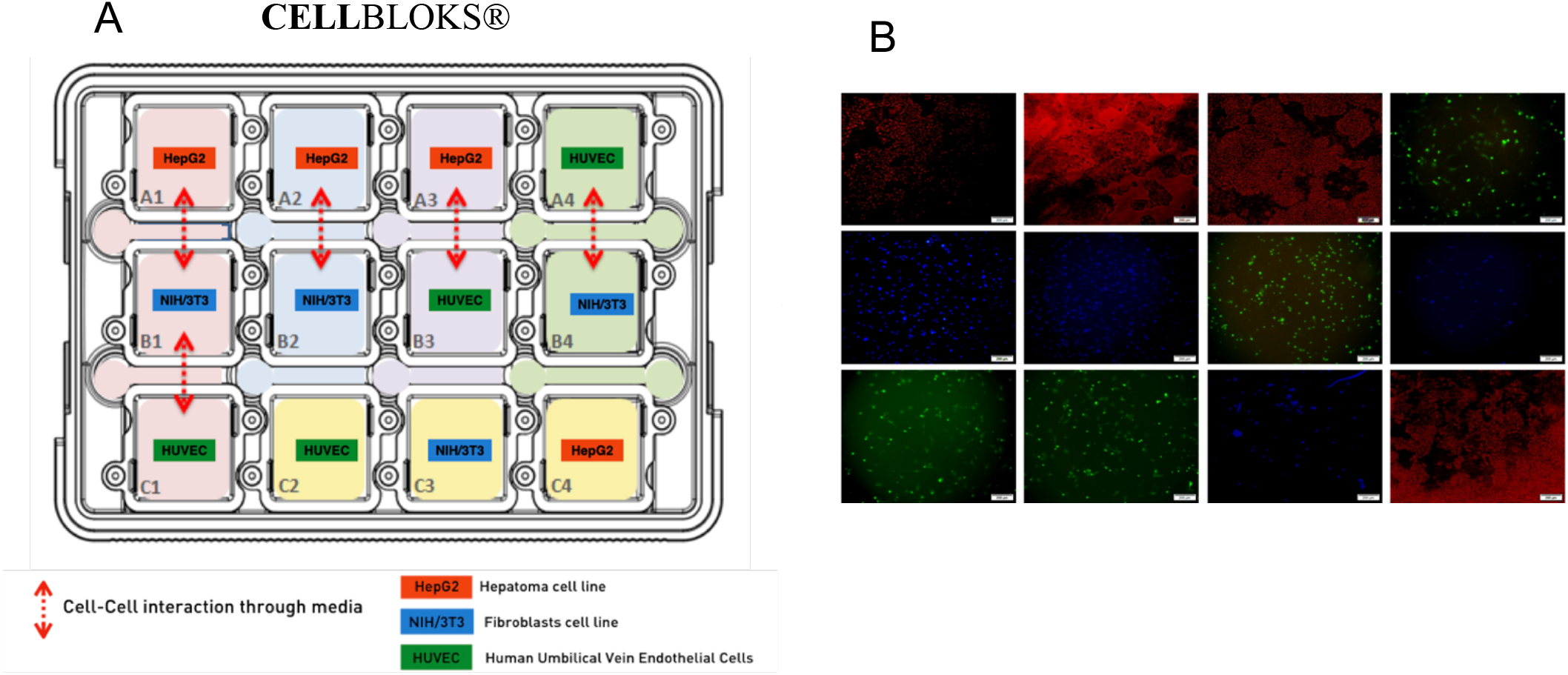
Liver model set-up on CELLBLOKS^®^ platform; (**A**) images of different cells in their respective cell growth blocks (**B**). CELLBLOKS^®^ platform co-cultures were set-up using Circulatory Blocks with 1.0 μm pore size PC membrane. Non-contact cell-cell interactions were tested in a tri-culture [A1--B1--C1], set of two combinations [A2--B2], [A3--B3] and [A4--C4] and in isolation to determine which cell-cell combinations produced optimal hepatic relevance. Each cell type was also grown in isolation at the same time in Blank Blocks in [C2], [C3] and (C4) compartments. Cells were imaged in day 5 after culture in their respective platform using an Olympus IX73 Inverted Microscope, magnification ×10: HepG2 cells (red), HUVECs (green) and NIH/3T3 cells (blue).

To prepare the co-culture blocks, 1 ml of cell suspension of 1×10^5^ cells/well for each cell line (HepG2, NIH-3T3 and HUVEC) were seeded into each interconnected cell blocks with HepG2 on the first row, NIH-3T3 on the second row and HUVEC on the third row with 3 ml of mixed growth medium (DMEM and EGM) added to the circulating channels around the blocks. For the monoculture, 1 ml of cell suspension of 5×10^4^ cells/well of each cell line was seeded separately on the isolated blocks of CELLBLOKS^®^ plate. Cell Viability, albumin, and cytochrome P450 was measured every 2 day for 14 days.

### Viability assay

Viability of the three cell lines (HepG2 cells, HUVECs and NIH-3T3) was measured simultaneously in the CELLBLOKS^®^ platform for up to 14 days (Figure 3). Cell viability was assessed using the Alamar Blue cell viability assay (Invitrogen Thermofisher™). A stock solution was prepared according to manufacturers to instructions, and diluted to a working (AB) solution in a Hank’s balanced salt solution (Sigma-Aldrich™) and kept at 37°C. After the incubation time, the culture media were removed, and the cells washed twice with PBS. To detriment viability of individual cell types in co-culture set-ups, 1ml of the diluted AB solution was added into each block and the plates were incubated at 37°C and 5% CO_2_ for 1 h. After incubation, aliquots of 100 μl from these blocks and wells were transferred into a 24-well plate for reading. The intensity was read by a Synergy H1 microplate reader from BioTek™ (Absorption 530; Emission 590) and analysed with Gen5 software. The cells were then washed with 1 ml of PBS and 1 ml of media was added. The plates were re-incubated at 37°C and 5% CO_2_.

**Figure 3.**
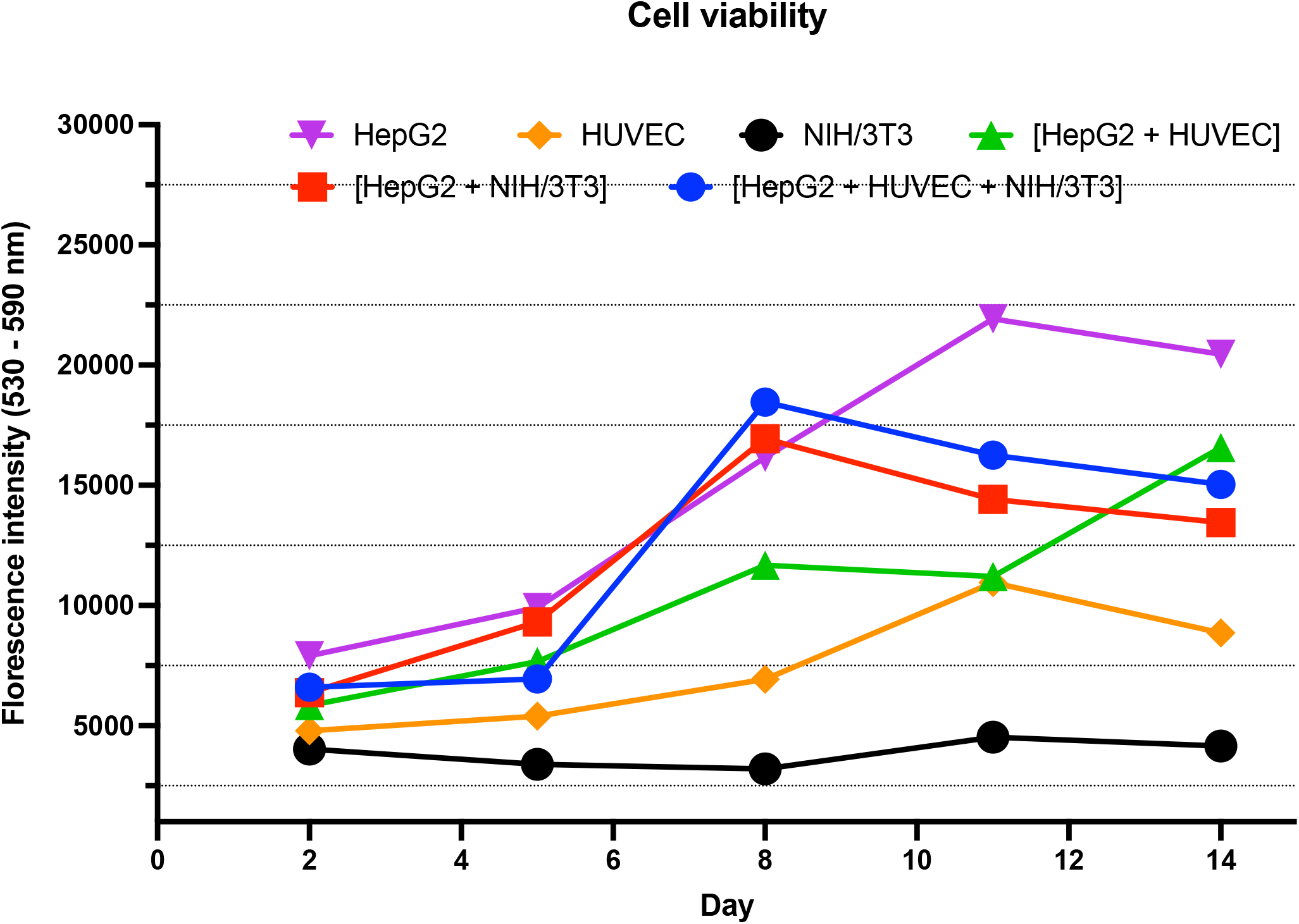
Viability of different cell types and co-culture set-ups on CELLBLOKS^®^ platform. Viability was measured in all cells types in monocultures: HepG2 cells, NIH-3T3 cells and HUVECs; as well as in HepG2 cells co-cultured with NIH-3T3 cells [HepG2 + NIH/3T3], HepG2 cells co-cultured with HUVEC [HepG2 + HUVEC] and in tri-culture [HepG2 + HUVEC + NIH/3T3] (n = 3).

### Imaging

HepG2 cells and HUVEC were stained with the Cell Tracker Red CMTPX (Thermofisher™) and the Cell Tracker Green CMFDA (Thermofisher™) respectively on day 0 of the experiment (Figure 2B). The cells were cultured in T25 flasks in appropriate media. To prepare the working solution, cell trackers were diluted in diméthylsulfoxyde (DMSO) (Sigma-Aldrich™) to a final concentration of 10 mM, and finally in serum-free media (SFM) (Gibco™) to a final concentration of 10 μM. Once the cells were confluent, the media was removed from the flasks, cells were washed twice with PBS, and the working solution of cell tracker was gently added (Red Cell Tracker to HepG2 cells and Green Cell Tracker to HUVEC). The cells were incubated at 37°C and 5% CO_2_ for 45 min. After the incubation, the solution from the cells was removed, the cells were washed twice with PBS, and trypsinized with 2 mL of trypsin (0.05%) for 7 min prior to being seeded in 12-well plate or CELLBLOKS^®^ platform according to the experimental plan. On day 1, 2 μM of Hoechst solution (Sigma-Aldrich™) was added to each well. The cells were imaged with an Olympus IX73 Inverted Microscope (Olympus™) using magnification ×10, with Olympus CellSens standard software. Images were processed with ImageJ software.

### Albumin assay

Albumin production from each cell growth block containing HepG2 cells was determined using the Bromocresol Green (BCG) Albumin assay kit (MAK124, Sigma-Aldrich™). Albumin standard curves were first calculated according to vendor’s protocol. Cells were scraped from each block into 1-ml separate centrifuge tubes and counted using a haemocytometer. For each condition cells were lysed in 100 μl of cold lyses buffer for 60 min at 4°C. The cell solution is centrifuged at 13000 g for 10 min at 4°C to remove insoluble material. In each well of a 96-well plate, 10 μl of sample supernatant (or a standard) and 200 μl of albumin reagent are added according to the supplier’s indication. The absorbance was measured with a Synergy H1 microplate reader from BioTek™ and analysed with Gen5 software. The assay is conducted in triplicate at different time points: day 1, day 4, day 8 and day 12 of culture.

### Urea assay

The biosynthetic capabilities of HepG2 cells were assessed using a Urea assay kit (MAK006, Sigma-Aldrich™). A standard curve was measured, and urea production was measured according to the supplier’s instructions. Cells were first scraped form each block into 1 ml separate centrifuge tubes and counted using a haemocytometer. Then for each condition HepG2 cells were lysed in 100 μl of cold lyses buffer for 60 min at 4°C. The cell solution was centrifuged at 13000 g for 10 min at 4°C to remove insoluble material. In each well of a 96-well plate, 50 μl of sample supernatant (or standard) and 50 μl of enzyme reaction mixes (peroxidase substrate, enzyme mix, developer and converting enzyme) are added according to the supplier’s indication.

The absorbance was measured with a Synergy H1 microplate reader from BioTek™ and analysed with Gen5 software. The assay is conducted in triplicate at different time points: day 1, day 4, day 8 and day 12 of culture.

### Cytochrome P450 assay

CYP3A4 expression in HepG2 cells was measured with a P450-Glo^TM^ assay (V9001 Luceferin-IPA, Promega™). Cells were scraped from each block into 1 ml PBS tubes and counted using a haemocytometer. Cells were then placed in 96-well opaque plates and the assay was performed according to the supplier’s instructions. Luminescence was measured with a Synergy H1 microplate reader (BioTek™) and analysed with Gen5 software. The assay was conducted in triplicate at different time points: day 1, day 4, day 8 and day 12 of culture.

### Tamoxifen toxicity on triculture *versus* monoculture

All dilutions were prepared using sterile culture media in a sterile culture hood. To prepare the triculture blocks, 1 ml cell suspension of 5×10^4^ cells/well for each cell line (HepG2, NIH-3T3 and HUVEC) were seeded into the interconnected Circulator blocks™ with HepG2 on the first row, NIH-3T3 on the second row and HUVEC cells on the third row with 3 ml mixed growth medium (DMEM and EGM) added to the circulating tunnels around the blocks; these were incubated for 24 h. For the monoculture, 1 ml cell suspension of 5×10^4^ cells/well of only the HepG2 cell line was seeded on the isolated Blank blocks™ of a CELLBLOKS^®^ platform. Treatment with different tamoxifen concentrations (0.1 μM, 1 μM, 10 μM, 50 μM, 100 μM) for each cell line using DMSO as control, 60 μL of DMSO was added to the first row (HepG2, NIH-3T3 and HUVEC) by adding 10 μl to each block and 30 μl in the circulating medium. For the drug treatment the same sequence was followed (60 μl of each drug concentration was distributed between blocks and circulating medium), then incubated for 24 h. Cytotoxicity of tamoxifen was measured on HepG2 cells using CellTiter-Glo^®^ Luminescent ATP Cell Viability Assay (G7570, Promega) and the assay was performed according to the manufacturer’s protocol. Briefly, the plates with its contents were equilibrated to room temperature for approximately 30 min, then media was removed from each block and replaced with 200 μl fresh media and 200 μl of CellTiter-Glo® reagent (CellTiter-Glo® Buffer plus CellTiter-Glo® lyophilized substrate) and incubated at room temperature for 10 min. Luminescence was recorded using a microtiter plate reader. Cell viability was expressed in percentages in comparison to control (DMSO treated) (*n*=12).

### Statistical analysis

All statistical analysis were performed using GraphPad Prism (V9.3.1). Analysis of statistical significance comparing Urea, Albumin and CY3A4 in Monocultures *versus* Co-culture and Tri-culture conditions were calculated using two-way multiple comparisons ANOVA. A *P*-value of \**P* ≤0.05 was set as a threshold for statatistically significant results. IC50 values of tamoxifen dose-respose curves were measured using non-linear four parameter variable slope model, Log (inhibitor) *versus* Response.

## RESULTS

Cell viability was determined in cell growth blocks for a period of 14 days culture and shows that the platform supports cell growth with different monocultures and co-cultures that vary in their growth rate and pattern (Figure 3). In general, HepG2 monocultures grow at higher rate compared to both HUVEC and NIH-3T3 monocultures and this is apparent in the 2-week duration of cell growth tests. HepG2 cell viability alone shows a gradual temporal viability increase from day 2 to day 11, after which they start to decline (Figure 3). When HepG2 cells are connected in co-culture with HUVECs, viability increases continuously from day 2 to day 8 and decreases slowly after day 8. The viability curve of HepG2 cells connected with NIH-3T3 cells has the same pattern; cells grow rapidly between day 5 and day 8 of culture, and viability decreases slowly after day 8. Finally, HepG2 cells growth in tri-culture with HUVECs and NIH-3T3 cells is initially slow during the first 5 days of co-culture and then cell viability increases rapidly until day 8 and decreases slowly in a similar pattern to other coculture combinations (Figure 3).

Imaging of cells through the platform where each cell type was pre-labelled with different cell tracker (HepG2 cells in red, HUVEC cells in green and NIH-3T3 cells with the DNA intercalating agent Hoechst in blue) before seeding. Figure 2B shows live imaging of cells compartments within each cell growth block where cells were labelled directly without being disturbed. Tracking of cells with time indicated that cells seeded independently remain in their blocks and are unable to pass through membrane to other cells’ compartments. This allows each cell type to be studied individually without cross-contamination.

Following assessment of viability and imaging of cells in the platform, levels of urea, albumin and CYP3A4 activity were measured in HepG2 hepatocytes at different time points to determine hepatic relevance effected by different cell combinations (Figure 4–6). Urea production/cell was generally higher in days 1 to 4 of culture in all conditions but decreased in a time-related fashion after day 4 with the lowest levels of expression noted on day 12 (Figure 4). At day 1, urea levels remain similar in monocultures as well as other co-culture conditions. However, at day 4 there is a significant effect of either cell co-cultures on urea produced by HepG2 cells, where urea expression is doubled by the presence of HUVEC cells (~40 pg/cell) and tripled by the presence of NIH-3T3 fibroblasts (~60 pg/cell) co-cultures, respectively. However, in tri-cultures (HepG2+HUVEC+NIH/3TR) urea level of ~30 pg/cell is noted, although this is not statistically significant at *P* =0.05.

**Figure 4.**
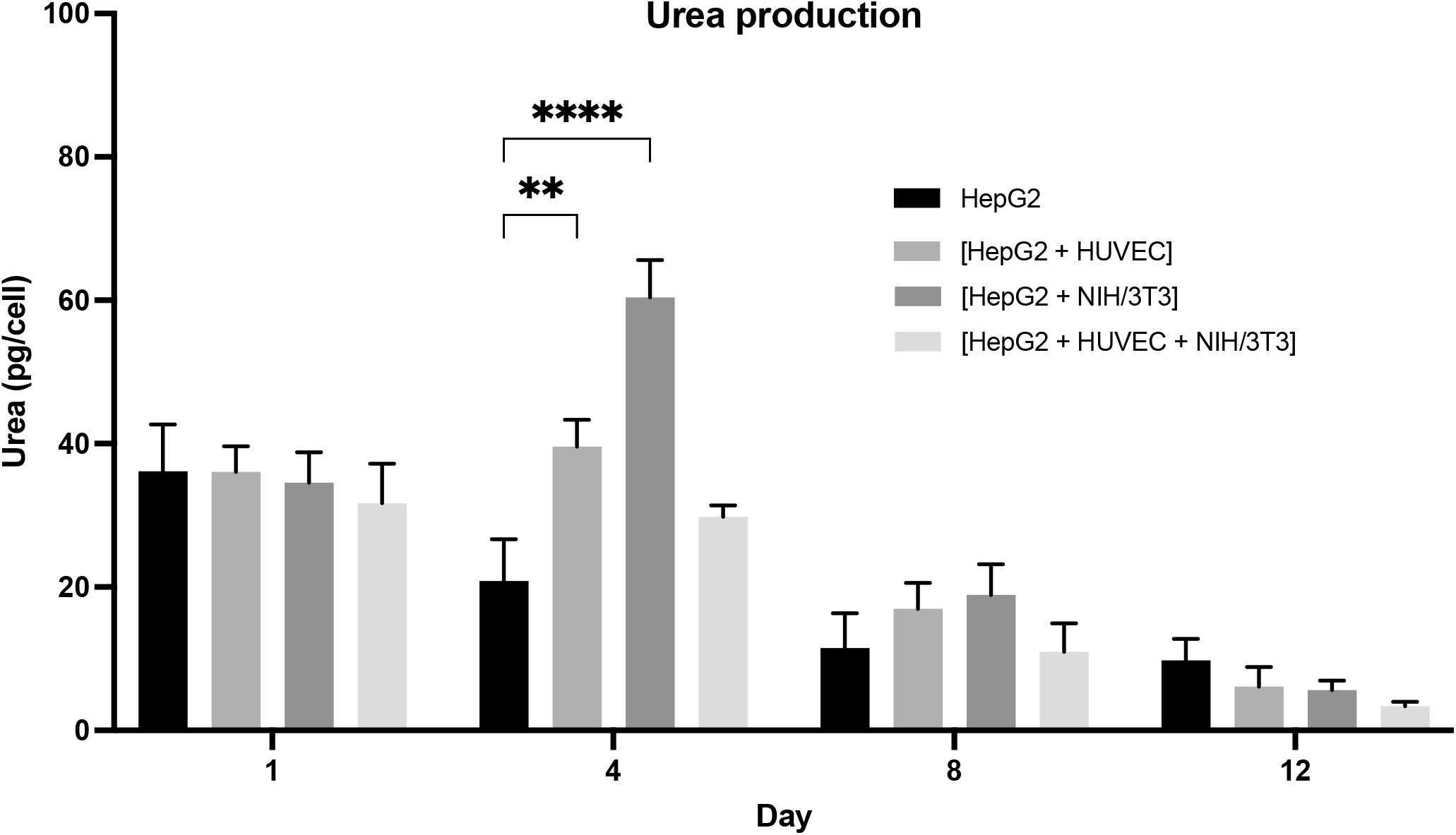
Urea Production in HepG2 cells in different cell co-culture and tri-culture condtions. Urea was measured in HepG2 cells alone (HepG2), HepG2 cells co-cultured with NIH-3T3 cells [HepG2 + NIH/3T3], HepG2 cells co-cultured with HUVECs [HepG2 + HUVEC] and in tri-culture [HepG2 + HUVEC + NIH/3T3] (*n*=3, ± SEM). Urea production in HepG2 monocultres was compared to co-cultures and tri-cultures repectively using two-way ANOVA in GraphPad Prism 9 (\**p* ≤0.05, \*\**p* ≤0.01, \*\*\**p* ≤0.001).

Similarly, albumin levels are at their highest in the first 4 days of culture and then their levels fall away after day 5; this is apparent in all culture conditions (Figure 5). Co-culture is noted to induce a significant increase in albumin levels compared to HepG2 cells cultured independently. However, co-culture does not appear to improve albumin production in the first 24 h of incubation, where a reduction of albumin levels is observed in HepG2 + NIH-3T3 co-cultures and tri-cultures of HepG2 + HUVEC + NIH/3T3. The effect of co-cultures in HepG2 albumin production is most noticeable at day 4 of culture where albumin levels are markedly reduced in HepG2 monocultures (from 25.6 pg/cell down to 6.2 pg/cell) but in cocultures levels range from 9.3 pg/cell to 23 pg/cell, it being noted that the presence of NIH-3T3 cells give the most positive response. In all cell growth conditions, CYP activity remains high in the first 24 h of culture in all conditions but reduces significantly after this point (Figure 6). All co-cultures conditions significantly improve CYP3A4 expression following 24 h of co-culture; HUVEC and NIH-3T3 cells stimulate HepG2s to double CYP3A4 production from 21.6 ng/cell to 44.3 pg/cell and 45.9 pg/cell respectively. However, in tri-culture (HepG2 + HUVEC + NIH/3T3) levels of CYP3A4 expression are tripled to levels of 67.1 pg/cell.

**Figure 5.**
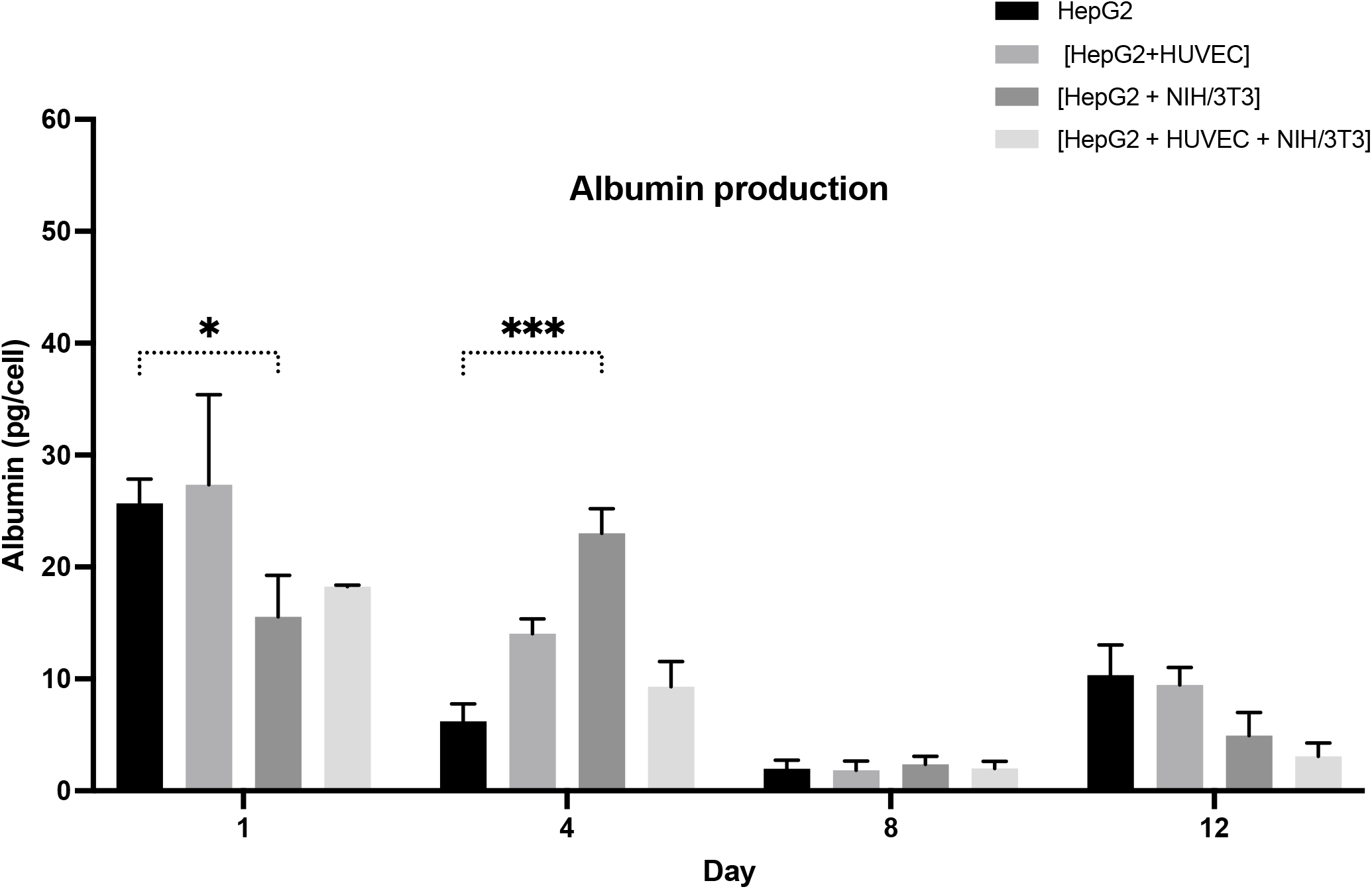
Albumin production in HepG2 cells in different cell co-culture and tri-culture condtions. Urea was measured in HepG2 cells alone (HepG2), HepG2 cells co-cultured with NIH-3T3 cells [HepG2 + NIH/3T3], HepG2 cells co-cultured with HUVEC [HepG2 + HUVEC] and in tri-culture [HepG2 + HUVEC + NIH/3T3] (*n*=3, ± SEM). Albumin levels in HepG2 monocultres was compared to co-cultures and tri-cultures repectively using two-way ANOVA in GraphPad Prism 9 (\**p* ≤0.05, \*\**p* ≤0.01, \*\*\**p* ≤0.001).

**Figure 6.**
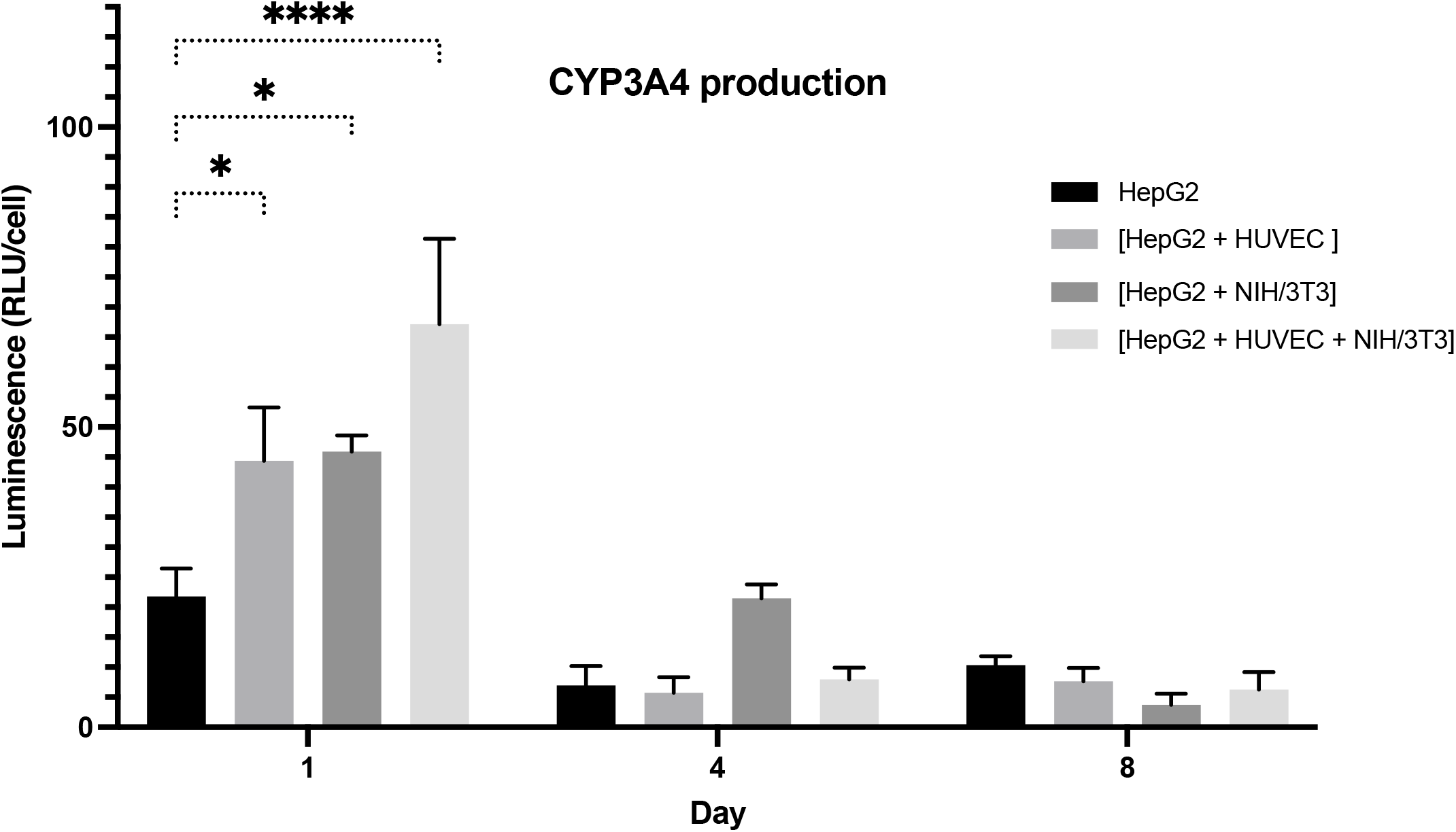
CYP3A4 production in HepG2 cells in different cell co-culture and tri-culture condtions. CYP3A4 was measured in HepG2 cells alone (HepG2), HepG2 cells co-cultured with NIH-3T3 cells [HepG2 + NIH/3T3], HepG2 cells co-cultured with HUVEC [HepG2 + HUVEC] and in tri-culture [HepG2 + HUVEC + NIH/3T3] (*n*=3, ± SEM). CYP3A4 levels in HepG2 monocultres was compared to co-cultures and tricultures repectively using two-way ANOVA in GraphPad Prism 9 (\**p* ≤0.05, \*\**p* ≤0.01, \*\*\**p* ≤0.001).

Tamoxifen induced dose-response effects in both HepG2 cells cultured alone and in HepG2 cells cultured in combination with HUVEC and NIH/3T3 cells, with lowest observed effects at 0.1 μM and more than 90% cell death at >50 μM concentrations (Figure 7). However, the dose response curve varied with each cell model. HepG2 cells in triculture exhibit higher sensitivity to tamoxifen treatment at lower concentrations <5 μM compared to HepG2 monocultures. Additionally, the IC50 value for HepG2 monocultures are double (16.40 *versus* 8.96) that of HepG2 cells in tri-culture, indicating a higher sensitivity to tamoxifen toxicity.

## Discussion

The main objective of this study is to demonstrate the capability of a new organ-on-a-chip platform (CELLBLOKS^®^) that can be used to create organotypic cell culture conditions in standard laboratory settings and test complex cell-cell interactions that give optimal biological relevance. We used the platform’s “plug and play” approach to model the liver *in-vitro* so that it can predict drug-induced hepatotoxic effects in humans more accurately during pre-clinical testing stages. The liver is a complex organ that involves the interplay of multiple cell types including hepatocytes, endothelial, fibroblast, bile duct epithelial and Kupffer cells (McCuskey, 2012). CELLBLOKS^®^ allows one to explore how heterogeneous interactions of different liver cell types drive hepatic relevance. The platform enables precise control of ratios between different cell types allowing various cell-cell interaction studies as well as facilitating the relative proximity between different cell types relevant to *in-vivo* locations. In addition, each cell population is grown in different compartments allowing cell proliferation in a specific required culture condition, which may be different from another cell type. Once the required cellular growth of each cell type is achieved, the cell growth blocks can be plugged in the platform for inoculating the co-culture. This particularly feature is useful with cells sensitive to media changes and having different growth rates.

Other recent methods for modelling liver *in-vitro* include microfluid-chip approaches that also take into account-controlled perfusion in the cultures. Hepatocytes in the liver are not subjected to direct blood flow and are protected from flow-induced shear stress by endothelial cells that fenestrate in the capillary sinusoid wall allowing exchange of O_2_, CO_2_, and metabolites (Allen & Bhatia, 2003). Although, the introduction of flow can improve nutrient supply and induce O_2_ zonation, hepatocytes might also be subject to damage if the flow rates are too high (Kietzmann *et al*., 1996; Tanaka *et al*., 2006).

In CELLBLOKS^®^ liver model, the HepG2 cells are divided by a thin permeable membrane that allows metabolite and gas exchange between cells and the culture medium *via* passive diffusion and the flow is provided using the CELLBLOKS^®^. All the three cell types (HepG2, fibroblasts and endothelial cells) retain high levels of cell viability, possess excellent cellular morphology, and exhibit significantly enhanced liver functions compared to their counterparts in standard monocultures. The data suggests that proliferation of HepG2 cells has enhanced due to cellular interaction with other cell types (endothelial and fibroblast) and the flow system provided by the CELLBLOK^®^ platform. Further to that, the liver function in coculture of all three cell types is significantly improved compared to monoculture hepatocytes, this includes albumin protein production, CYP450 expression and urea synthesis; this highlights the need for their inclusion in liver modelling to mimic physiological relevant human liver. Though the CYP activity of HepG2 cells was high in the first 4 days of culture a subsequent decline was observed, which is in line with Duthie and Collins’ study that reported that reduced glutathione content dramatically increases at 24 h of HepG2 cell culture and declined after one day of culture when cells approach confluence (Duthie & Collins, 1997). Although HepG2 cells express low amounts of CYP compared to HepRG and primary hepatocytes they are still used in different drug screening programs due to being well-established, widely available and extensively studied compared to other cells (Bale *et al*., 2014; Gebhardt *et al*., 2003; Gomez-Lechon *et al*., 2008; Gómez-Lechón *et al*., 2014). Compared to conventional mono-cultured HepG2 cells, co-cultures stimulated an increase of CYP3A4 metabolic activity by up to three times indicating an improved *in vitro* liver model. Given that CYP450 enzymes are important in drug metabolism, drug-drug interactions, and are commonly used to measure drug induced cellular responses (McDonnell & Dang, 2013), the tri-culture model (HepG2 + HUVEC +NIH/3T3) was selected to examine tamoxifen toxicity as this model exhibited the highest CYP3A4 compared to other co-culture and monoculture set-ups.

As HepG2 cells are derived from cancers, this would highlight the need for additional use of primary hepatocytes to further validate DILI predictions. CELLBLOKS^®^ platform allows one to perform experiments on any seeded cell type separately. It means that you can unplug the required cell type from the system any time during or after the experiment to study the effect separately. For example, we have studied hepatocyte metabolism in the HepG2 cell only from the pool of fibroblast and endothelial cells. Additionally, each cell type can be further analysed separately following a combined exposure to similar treatment regimes. This is particularly useful in understanding chemical mode of action at organ level rather than just the individual cell type. This feature is possible in the CELLBLOKS^®^ platform but not feasibly in randomly mixed co-culture models. For example, using our technology we analysed for liver function (*i.e*., albumin production, urea excretion and CYP activity) separately after coculturing the three cells for 14 days. This is important when albumin is also expressed in extra-hepatic tissues (Shamay *et al*., 2005) and urea is produced by HUVECs *via* arginase activity (Bachetti et al., 2004) and that may reduce the bias induced by the mix of two or three cell lines in the wells. In addition, tamoxifen toxicity when compared in monoculture and triculture (HepG2 separated analysis after culture) appeared to be increased in tri-cultured hepatocytes compared to mono-cultured hepatocytes indicating of increased sensitivity to toxic insult with this model.

The CELLBLOKS^®^ platform is a new promising cell culture device used for modelling cell-to-cell and organ-to-organ interactions. In this study, we have concentrated on cell-to-cell interactions to develop a liver model but the design of interconnected chambers allow organ-to-organ communication. The 3D organ/organoids (*e.g*., gastro-intestinal tract, liver) are grown separately and then plugged together in combined culture conditions. For instance, Barrier blocks™ containing selective membrane at the bottom can be used to create barrier functions (*i.e*., gastro-intestinal tract or blood brain barrier), whereas Circulatory blocks™ are applied to simulate for non-barrier organ functions. The Blank blocks™ are used in isolating cultures as controls to study their function compared to connected interacting cells.

Furthermore, seeding the cells in independent compartments makes experiments easier while maintaining communication *via* soluble factors excretion. When tri-cultures or co-cultures are carried out, assays can be performed only with the cell line of interest. Another advantage is the compatibility of the plate with readout equipment such as microscopes and plate readers as well automated handling applied in high-throughput screening (HTS). Moreover, the “plug & play” deign makes it easy to use, like a conventional well plate; its utilization is easier than complex microfluidic systems.

The CELLBLOKS^®^ is designed to explore crosstalk between tissues and cells in separate cultures. In this system, different types of cells can be cultured individually but connected through the flow of the medium. This enables each culture to be addressed and interrogated individually. While conventional co-cultures are useful for the study and optimization of cell function, they are not a suitable model for investigating interactions between cell types arising from different tissues. In this sense, the CELLBLOKS^®^ platform represents a more physiological relevant model. Herein, hepatocytes’ functionality when combined with endothelial cells and fibroblast in both monocultures, co-cultures and tri cultures was explored, which highlighted that cell-cell interactions produce most biological relevance. Connected cultures enhance albumin synthesis, urea production and CYP metabolic activity in hepatocytes compared to non-connected monocultures. Therefore, as demonstrated here, the connected culture in the CELLBLOKS^®^ system combines the dynamic stimulus of flow with cell crosstalk through soluble ligands so that the unit production of albumin, CYP3A4 and urea is greater than in monocultures.

In summary, we have developed a novel device that can be used to culture cells and produce cellular models with optimal organ specific biological relevance. We have demonstrated the application of this technology for the human liver model and have shown that cellular performance can be significantly enhanced. *In-vitro* systems that enable cells to grow in a manner more closely resembling their native counterparts will result in the development of assays that provide more accurate data about cell function which in turn will contribute to improving the efficiency of research and development. We demonstrate that CELLBLOKS^®^ interactive “plug and play” approach allows one to systematically test the interactions of multiple cell types simultaneously in one platform that helps to unravel complex cell-cell interactions that drive biological relevance. In addition, we hypotheses that multiple cell type engineering of liver model in such non-contact setup is more advantageous for screening DILI issues compared to randomly mixed co-cultures both practically and as a predictive assay.

## Acknowledgements

We wish to thank Innovate UK for funding this project (Grants 103006, 104432). CELLBLOKS^®^ is a patented technology and consists of family of patents including UK (Granted; GB2553074B), US (Pending: 16/075,136), PCT (Pending: PCT/GB2017/05) and Europe (Pending; EP:1713365.9).

